# Just the Two of Us? A Family of *Pseudomonas* Megaplasmids Offers a Rare Glimpse Into the Evolution of Large Mobile Elements

**DOI:** 10.1101/385575

**Authors:** Brian A. Smith, Courtney Leligdon, David A. Baltrus

## Abstract

Pseudomonads are ubiquitous group of environmental proteobacteria, well known for their roles in biogeochemical cycling, in the breakdown of xenobiotic materials, as plant growth promoters, and as pathogens of a variety of host organisms. We have previously identified a large megaplasmid present within one isolate the plant pathogen *Pseudomonas syringae*, and here we report that a second member of this megaplasmid family is found within an environmental Pseudomonad isolate most closely related to *P. putida.* Many of the shared genes are involved in critical cellular processes like replication, transcription, translation, and DNA repair. We argue that presence of these shared pathways sheds new light on discussions about the types of genes that undergo horizontal gene transfer (i.e. the complexity hypothesis) as well as the evolution of pangenomes. Furthermore, although both megaplasmids display a high level of synteny, genes that are shared differ by over 30% on average at the amino acid level. This combination of conservation in gene order despite divergence in gene sequence suggests that this Pseudomonad megaplasmid family is relatively old, that gene order is under strong selection within this family, and that there are likely many more members of this megaplasmid family waiting to be found in nature.

## Introduction

Horizontal Gene Transfer (HGT) of megaplasmids can rapidly create dramatic phenotypic differences between otherwise closely related bacterial strains, with potential for over a thousand genes to be gained by a strain in a single event. Although there have been numerous attempts to identify overarching themes for the evolutionary effects of HGT based on types of genes and pathways transferred, such efforts have often neglected to incorporate intrinsic characteristics of megaplasmids ^1-6^. Furthermore, since secondary replicons are prone to rapid reshuffling of gene order as well as extensive gains and losses of loci, it has traditionally been challenging to analyze past evolutionary dynamics to understand overall historical pressures acting on this class of mobile elements^7-14^. More thorough investigation of evolutionary dynamics within relatively large plasmid families could therefore provide new viewpoints into the evolutionary effects of gene transfer and may also enable broader generalizations about selective forces driving the composition and overall structure of megaplasmids, chromids, and second chromosomes.

Megaplasmids are generally characterized as low copy extrachromosomal replicons >350kb in size and which are dispensable to the bacterial cell under a subset of conditions^14^. As with many plasmid families, they have often been identified because they impart beneficial phenotypes such as resistance to antimicrobial compounds or introduce novel catabolic pathways into host cells^14^. Given their size and gene content, it is possible that megaplasmids possess greater potential for generating evolutionary costs than smaller plasmids when transferred to naive hosts^15,16^.

However, efforts to identify shared genes and pathways across megaplasmids and to use this information to make predictions about potential systems level conflicts have been hampered by poor sampling across novel megaplasmid families ^14^. Identification of additional examples can help to fill this gap in current knowledge and may uncover new evolutionary trends that govern megaplasmid-chromosomal interactions.

Consideration of megaplasmids may uniquely inform general discussions about evolutionary effects HGT in ways that have been overlooked by analyses focusing simply on distributions of genes and pathways maintained after transfers and without considering timing of HGT or linkage between loci. For instance, many of the earliest discussions concerning evolutionary constrains of HGT, the so called “complexity hypothesis”, found that loci associated with “more complex” cellular processes undergo lower rates of transfer than other genes^4,17^. Interpretations of these patterns have changed through time with examples of horizontally acquired informational genes, suggesting that it is actually the shape of protein interaction networks that are critical for maintenance after gene transfer. ^4,18-22^. However, larger mobile elements like megaplasmids can contain genes encoding proteins and pathways that could be classified as “complex” (i.e. proteins involved in translation) and which have the potential to interact with numerous chromosomally encoded pathways^14,23^. Likewise, a variety of recent papers have focused on selective forces (or lack thereof) governing microbial pangenomes^1-3,24^. These discussions have largely focused on population level distributions for single genes that compose a pangenome, but by their nature intrinsically fail to consider linkage of genes on plasmids that are frequently acquired and lost. While many genes within the pangenome may indeed be ‘adaptive’, such a viewpoint overlooks the idea that no single gene need be adaptive for the bacterial cell if selection acts at the level of plasmid transfer and hundreds of genes that could be linked to that process. More thorough characterization of multiple megaplasmid families and identification of new megaplasmids will enable identification of the types of genes and pathways canonically associated with these large vectors and patterns that emerge can be incorporated into greater discussions of the general role of HGT across bacterial species.

Previously we have described a megaplasmid, pMPPla0107, found within one isolate of the phytopathogen *Pseudomonas syringae^23^.* pMPPla107 is self-transmissible across the *Pseudomonas* phylogeny, harbors numerous loci that could be annotated as “housekeeping” genes, and it is stably maintained within recipient cells^23,25,26^. We have also demonstrated that acquisition of this megaplasmid through HGT also imparts significant phenotypic costs to recipient cells, likely mediated by detrimental interactions between chromosomal and plasmid encoded proteins^26^. It is unclear if pMPPla107 is the only megaplasmid of its kind and size with conjugation abilities across an entire genus of bacteria or if it is a member of a larger family of secondary replicons.

Here we identify a new megaplasmid, pBASL58, related to pMPPla107 and use molecular and computational approaches to characterize this megaplasmid family more broadly. We show that, while these megaplasmids are similar in size, genetic structure, nucleotide bias, and functionality, there is a high level of divergence across shared orthologous gene groups, a dissimilar cargo region, and differing CRISPR loci. This overall level of divergence suggests that both members of this megaplasmid family have been independently evolving for a relatively long period of time. Even more, we find that these divergent orthologous pathways demonstrate high levels of synteny in the context of overall plasmid structure; suggesting that conservation of gene orientation and order is under relatively strong selective pressures. Lastly, characterization of these plasmids allows for a chance to emphasize that pathways found on both megaplasmids are likely involved in important cellular processes like nucleotide synthesis and DNA replication, which highlights new discussion points to add to the complexity hypothesis as well as the adaptive nature of pangenomes.

## Methods

### Identification of pBASL58

Initial BLASTp searches coupled with inspection of contigs from a draft genome assembly of *Pseudmonas* sp. Leaf58^27^ suggested that this strain could contain a megaplasmid related to pMPPla017, represented within contig Ga0102293_111within the draft genome assembly containing 18 contigs total (Genbank accession GCA_001422615.1). An isolate of *Pseudomonas* sp. Leaf58 was obtained from DSMZ (DSM-102683), and a single colony was picked from a culture from the freeze-dried ampule plated on unsupplemented LB media. To test for circularization of contig Ga0102293_111, primers (BAS 17-31) were designed to amplify off of the edges of the contig and overlap each other. An approximate 1.5kb size PCR product was amplified from an overnight culture of this strain grown in KB media, demonstrating circularization of this contig. Sanger sequencing of this PCR fragment demonstrated that the initial draft contig contained a missassembly, and so this sequence was corrected by hand, and the contig as reoriented and used in all analyses in this manuscript. Sequence of this contig can be found at Figshare (doi:10.6084/m9.figshare.6914033). For consistency of analyses throughout the manuscript we reannotated the megaplasmid sequence with Prokka v1.12 using default parameters, and this annotation can also be found at Figshare (doi:10.6084/m9.figshare.6914033).

### Genome Sequencing, assembly, and annotations of Pseudomonas sp. Leaf58

As further evidence of the existence of a megaplasmid in Leaf58, we generated a complete genome assembly for this strain (currently found at Figshare (doi:10.6084/m9.figshare.6914033). Genbank accession TBD). After revival from the Baltrus lab stock, a single colony *Pseudomonas* sp. Leaf58 was picked to an overnight culture in KB media, and grown in a shaking incubator at 27°C. After approximately 24 hours, DNA was extracted from this culture using a Promega Wizard kit. A rapid sequencing library was created using this DNA, and 169,316 reads (933,937,907 total bp, 5,515bp average read size) were generated on an R9.4 flowcell using a Rapid sequencing kit (SQK-RAD004). Additionally, 100bp paired end Illumina reads used to generate the original draft genome of this strain were downloaded from the SRA (Accession ERR1103815)^27^. A complete genome sequence for *Pseudomonas* sp. Leaf58 was generated by combining these short and long reads in Unicycler (version 0.4.4)^28^. This sequence consists of a single chromosome (5,432,868 bp) and the pBASL58 megaplasmid (904,253bp), both of which were circular according to Unicycler.

### Genome Sequencing, assembly, and annotations of Pla107

A single colony of the Baltrus lab stock of *Pseudomonas syringae* pv. *lachrymans* 107 (MAFF31015) was picked to an overnight culture in KB media, and grown in a shaking incubator at 27oC. After approximately 24 hours, DNA was extracted from this culture using a Promega Wizard kit. Illumina sequencing of was performed by MicrobesNG, and generated 2,771,213 250bp paired end reads (231 median read length after trimming, ~166x coverage of the genome) on an Illumina MiSeq. Assembly was performed using SPAdes v3.10 .1 with default parameters as well as through MicrobeNG’s bioinformatics pipeline, which matches the reads to the best reference using Kraken and maps reads back to that reference using BWA-MEM29. MicrobeNG also uses *de novo* assembly with SPAdes. pMPPla107 assembled completely from these reads, and this version of the megaplasmid sequence can be found Figshare (doi:10.6084/m9.figshare.6914033)and was used for all analyses throughout this manuscript. Gene annotation of this version of the megaplasmid sequence was performed with Prokka v1.12 using default parameters. This gene model used for all coding sequence analyses within the manuscript and can be found at Figshare (doi:10.6084/m9.figshare.6914033). We additionally generated long read sequences for Pla107 using a MinION from Oxford Nanopore. A rapid sequencing library was created from an independent genomic isolation of a derivative of Pla107, DBL328, which contains an integrated version of the pMTN1907 marker plasmid and which has been selected to for kanamycin resistance from this marker plasmid. As above, a single colony of this strain was picked to an overnight culture in KB media and DNA was extracted with a Promega Wizard kit. 15,461 reads (139,041,576 total bp, 8,993 average read size) were generated on an R9.4 flowcell using a Rapid sequencing kit (SQK-RAD004). A whole genome assembly was created by combining both MiSeq and MinION reads using Unicycler (version 0.4.4)^28^ with default parameters. This whole genome sequence consists of a circular chromosome (6,075,120 bp), pMPPla107 (971,889 bp, and sequence identical to the assembly from SPADES alone), and two other plasmids, pPla107-1 (62,136 bp) and pPla107-2 (40,720 bp). Three of these sequences (except pPla017-1) were complete and circular contigs according to Unicycler assembly. This assembly was used to update the Genbank version of this genome, and is found at accessions (CP031225, CP031226 CP031227, CP031228). Gene annotations in this Genbank file were generated by NCBI’s PGAAP pipeline^30^.

### Identifying Origins of Replication

To identify putative origins of replication for both megaplasmids, we used a modified GC skew script^31^ to scan the entirety of pMPPla107 and pBASL58 and combined this information with characterization of repetitive motifs that could represent *oriV* sites. GC skew and repetitive motifs suggest pMPPla107 and pBASL58 have predicted origins of replication within a similar genomic region near partitioning genes (Figure 2). Based on this information we oriented the sequences of pMPPla107 and pBASL58 to begin at the start codon of shared *parA-like* loci. We chose the parA-like locus as the starting point because it is shared by both sequences, is near the predicted origin of replication, and is predicted to be an important gene for plasmid partitioning.

**Figure 1:**
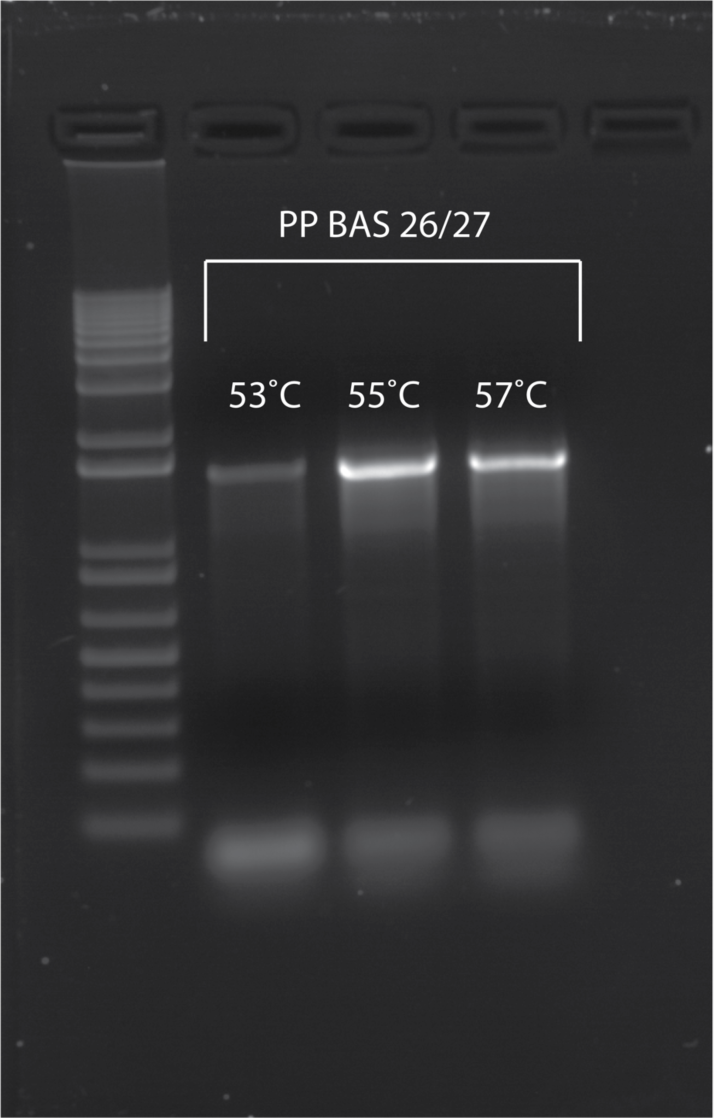
Confirmation of circular DNA molecule of Leaf58 megaplasmid. Primers designed to amplify the ends of the Leaf58 contig and disregarding the misassembled repeat region successfully amplified products of an expected size. Three annealing temperatures (53, 55, and 57°C) were used due to difficulties amplifying this region.

**Figure 2:**
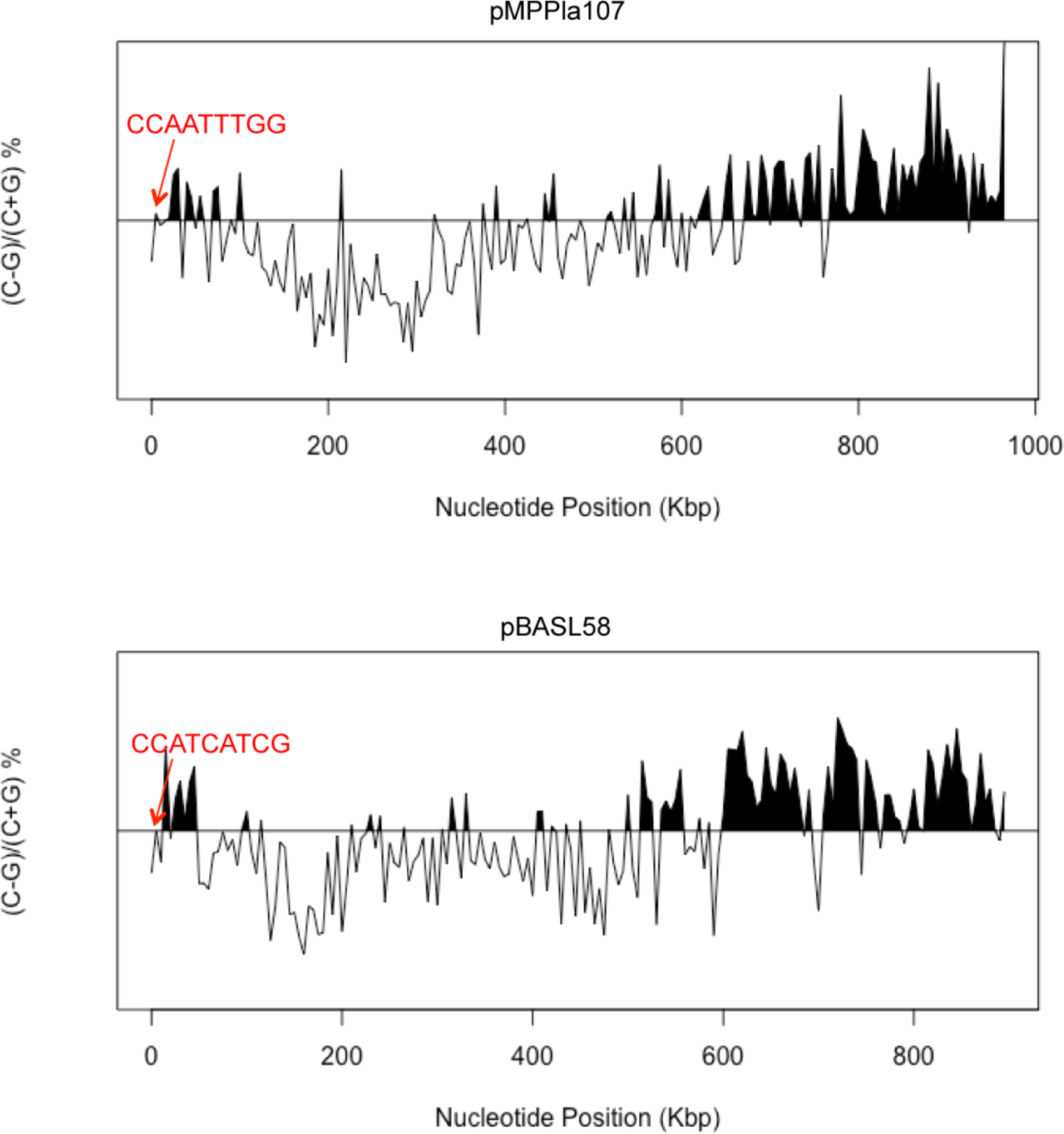
Predicted origins of replication occur within a similar region of pMPPla107 and pBASL58. GC skew was calculated and is a known predictor of origins of replication by a dramatic shift in GC content. Repetitive motifs were also calculated for areas near the predicted origin of replication as repetitive binding regions occur near replication sites. The most common motif is indicated in red.

### CRISPR Identification

CRISPR-Cas and repeat structure annotations were identified using both Prokka annotations and the web tool CRISPRCasFinder^32-34^

### Plasmid Comparisons With BLASTp and MAUVE

Amino acid sequence names were changed to numbers in an increasing order using the mod_protein_id.py script. We then used the BLAST 2.6.0+ package^35^. BLASTp parameters were altered to ensure only the top hit was returned and that there were zero overlapping hits. The BLAST command used was:

blastp -db [blastdb] -query [query_file] -culling_limit 1 -max_target_seqs 1 - max_hsps 1 -out [out_file] -outfmt 6

Data was extracted from the BLAST output at 40, 50, 60, and 70% identity cutoffs and plotted in R using ggplot2.

pMPPla107 and pBASL58 sequences were input into Progressive Mauve 2.3.1 to compare megaplasmid sequences within the software Geneious ^36^.

### Gene Mapping Visualization with Circa

The BLASTp output data mentioned above was altered in a format to comply with input to Circa using gff_info_extract.py followed by geneid_match.py. The parameters used to generate the Circa map and the Python scripts used to generate the data can be found at the https://github.com/basmith89/megaplasmid_compare.

### Tetranucleotide frequency Comparisons

We performed pairwise comparisons of tetranucleotide frequencies between chromosome sequences and secondary replicon sequences in an all by all method. Tetranucleotide frequencies were calculated with the calc.kmerfreq.pl script created by Mads Albertsen^37^ found at https://github.com/MadsAlbertsen/multi-metagenome. Output of this script was plotted using ggplot2 and R^2^ values were calculated in R.

### Functional Comparisons With KEGG, KASS, and UProC

We carried out two analyses utilizing the Kyoto Encyclopedia of Genes and Genomes (KEGG) database^38^. Amino acid sequences of coding regions predicted by Prokka were input into the protein sequence classification software, UProC^39^. UProC’s output is a list of KEGG IDs and counts. We designed a perl script, kegg_path_counter.pl, to extract these ID’s and counts and associated them with KEGG functional pathways. The script and ID key can be found at https://github.com/basmith89/megaplasmid_compare. These data were then plotted with the Plotly package in R.

Amino acid sequences output by Prokka were also run through a Python script to produce a list of gene annotations that both megaplasmids have in common https://github.com/basmith89/megaplasmid_compare. Amino acid sequences from genes on this shared list were then run through KASS (KEGG Automatic Annotation Server) to determine what pathways are shared by the megaplasmids^40^. Pathways maps were then condensed into one figure by hand.

## Results

### A new member of the pMPPla107 megaplasmid family

pMPPla107 was originally identified from an assembly using both 454 and 30bp Ilumina sequencing reads ^23^. However, due limitations of these early technologies, this assembly of pMPPla107 remained incomplete and consisted of linked scaffolds. We therefore utilized updated sequencing and assembly technologies to sequence the *P. syringae* genome containing pMPPla107, yielding a complete circular sequence for this megaplasmid (971,889bp compared to 963,598bp in original sequence) (Table 1). Additionally, multiple searches using protein sequences from pMPPla107 consistently yielded high quality matches to the scaffold Ga0102293_111 (referred to as pBASL58 from here on) from a public genome assembly of *Pseudomonas* sp. Leaf58. This strain was originally isolated as part of a project to thoroughly sample cultureable strains from the phyllosphere of Arabidopsis and is most closely related to *P. putida* strains^41^. We independently confirmed circularization (Figure 1) of this contig from Leaf 58, using both PCR and long read nanopore sequencing, definitively showing this contig was indeed a large megaplasmid separate from the chromosome.

**Table 1:**
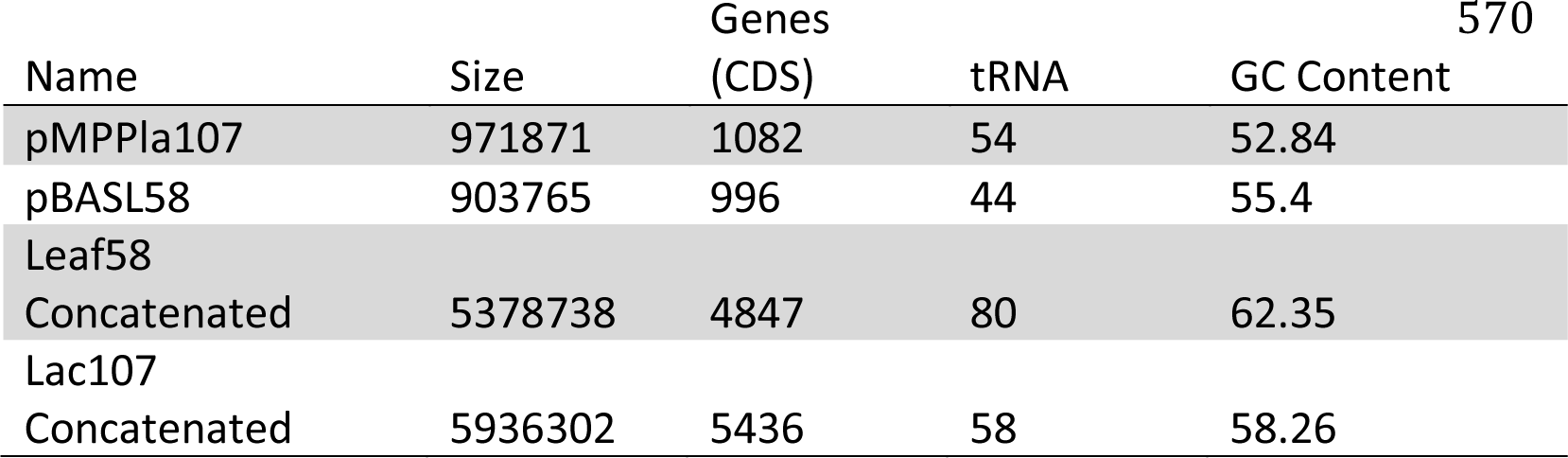
General features of *P. syringae* and *Pseudomonas* Leaf58 replicons have similarities. Size, coding regions, tRNAs, and GC content were calculated to understand the relationship between the four sequences on a broad scale. “Concatenated” indicates that all sequences from the assemblies expect either the pMPPla107 or pBASL58 replicons to their respective genome were concatenated together.

### Both megaplasmids contain numerous tRNA loci

The size, number of predicted genes, number of tRNAs, and GC content are highly similar between pMPPla107 and the pBASL58 (Table 1). Overall GC content was similar in Leaf58 and *P. syringae lac107*, and the GC content in both pMPPla107 and pBASL58 were lower than their respective chromosomal partners. pBASL58 and pMPPla107 contained 54 and 44 regions annotated as tRNA loci, respectively. pBASL58 encodes 20 unique tRNAs and pMPPla107 encodes 10, some of which were repetitive like tRNA-Glu(ttc) in pBASL58 occurring six times. When observing tRNA amino acid products, pMPPla107 encodes for 16/20 possible amino acids and pBASL58 encodes for 19/20 possible amino acids possibly indicating pBASL58 is less dependent on host tRNAs. In addition to the 16 amino acids produced by pMPPla107, pBASL58 is predicted to code for the ability to charge tRNAs with tryptophan, glutamate, and aspartate and both plasmids are missing any anticodons to produce histidine. These differences could suggest an amino acid preference for the maintenance or protein production of the plasmids.

### Identifying genomic similarities of pMPPla107 and the Leaf58 plasmid

Both megaplasmids within this new family are highly syntenic (Figure 3A, with Mauve alignment showing that 72.8% (707,677bp out of 971,871bp) of pMPPla107 aligns well with 71.6% (646,763bp out of 903,765bp) of pBASL58 (Supplemental Figure 1. The regions of highest similarity occur near the origin of replication. Despite overall high levels of synteny, there is a highly dissimilar region (approximately 300kb in size) occurring within the first half of the sequences and a ≈50kb inversion in the last half indicating these megaplasmids have also undergone structural diversification.

Even though both megaplasmids display high levels of synteny, preliminary comparisons of protein sequences suggested a relatively high level of divergence between orthologues shared by both megaplasmids (Figures 3B and C). The highest levels of average amino acid similarity (48.6%) occur near the predicted origin of replication where genes for plasmid replication, partitioning, and conjugation are common. Areas near the terminus still demonstrate strong synteny but have higher divergence in amino acid identity (≈38.2% similarity). These data suggest pMPPla107 and pBASL58 are structurally related to each other and share a common plasmid ancestor, but have experienced independent evolutionary pressures for long enough time for significant diversification to occur within shared protein sequences.

**Figure 3:**
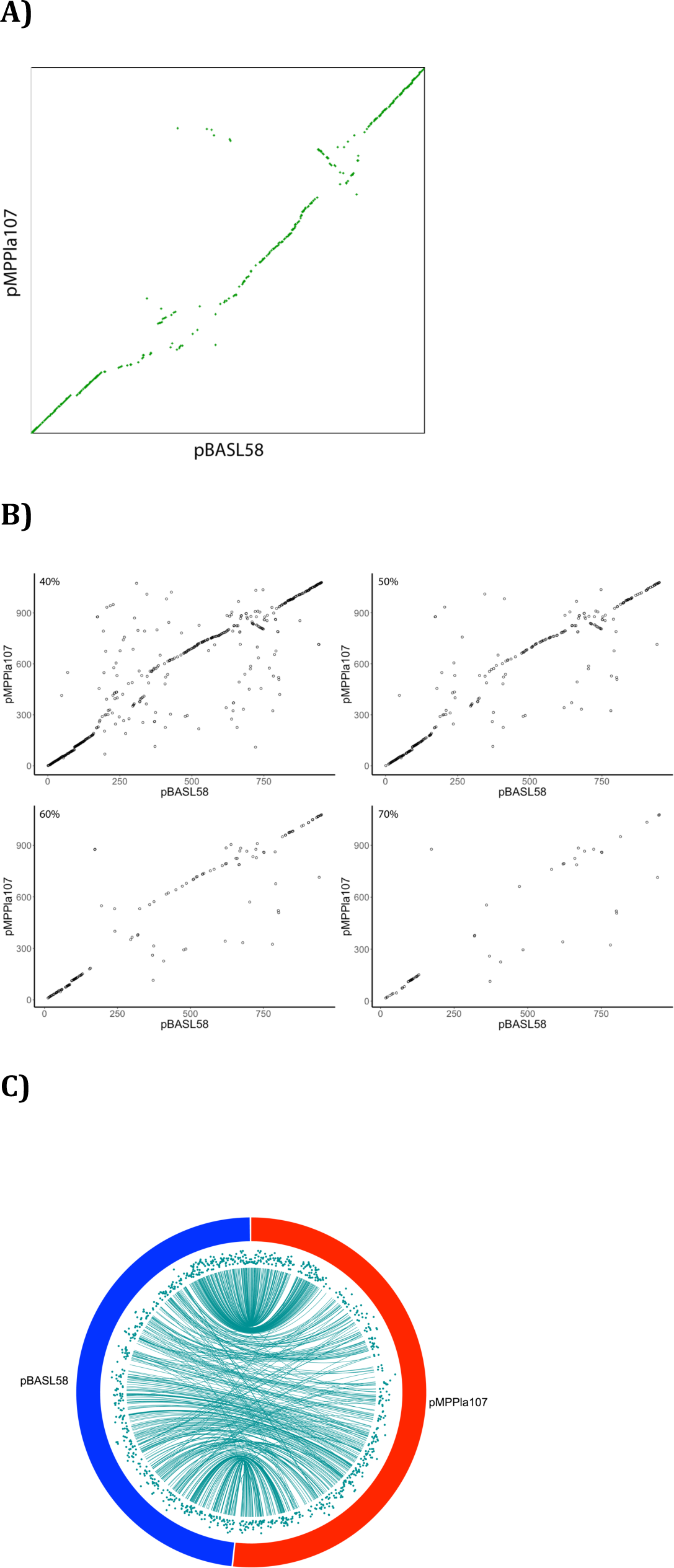
pMPPla107 and pBASL58 share synteny and demonstrate divergence on the amino acid level. **A)** SynMap output of pMPPla107 vs. pBASL58 sequences suggests highly syntenic megaplasmids. *X*-axis is pBASL58 gene order where *X*_I...N_ = gene_1...N_, and the *y*-axis is pMPPla107 gene order where *y*_1...N_ = gene_1...N-_ Completely syntenic sequences would be represented by y= 1x + b. **B-C)** BLAST data was used to plot pMPPla107 vs. pBASL58 synteny and amino acid divergence data together. **B)** The majority of syntenic genes have ≥50% sequence identity. BLAST data was plotted in gene order to mimic SynMap’s plot with amino acid sequence identity cutoffs at 40%, 50%, 60%, and 70%. The best hit for each pBASL58 gene against pMPPla107 is plotted. Each axis indicates gene position within the corresponding sequence. **C)** Circa plot using BLAST data indicates higher synteny near the origin, while areas near the terminus are less syntenic an experience more noise. Teal lines connect gene start position on pBASL58 to gene start position on pMPPla107. Teal scatter plots are amino acid sequence identity with 40% = 0 (bottom) and 100% = 100 (top)

To further gauge relationships between both megaplasmids and the chromosomes of their host strains, we compared tetranucleotide frequencies for each of these replicons ^42-44^. Pairwise comparisons demonstrated that pMPPla107/pBASL58 (R^2^ = 0.878) and the *P. syringae/Leaf58* (R^2^ = 0.889) chromosomes are most similar in frequencies (Figure 4). All remaining pairwise comparisons reported R^2^ values less than 0.780. pMPPla107 shows the greatest differences in tetranucleotide frequencies when compared to both the *P. syringae* and the Leaf58 chromosomes with R^2^ values of 0.524 and 0.393 respectively. pBASL58 shares slightly more similar frequency preferences indicative of R^2^ values of 0.780 and 0.695, to *P. syringae* and Leaf58 chromosomes respectively. This data suggest that mutational biases affecting these secondary replicons are most similar to each other, which suggests that they have not been replicating within these host strains long enough to be subject to amelioration.

**Figure 4:**
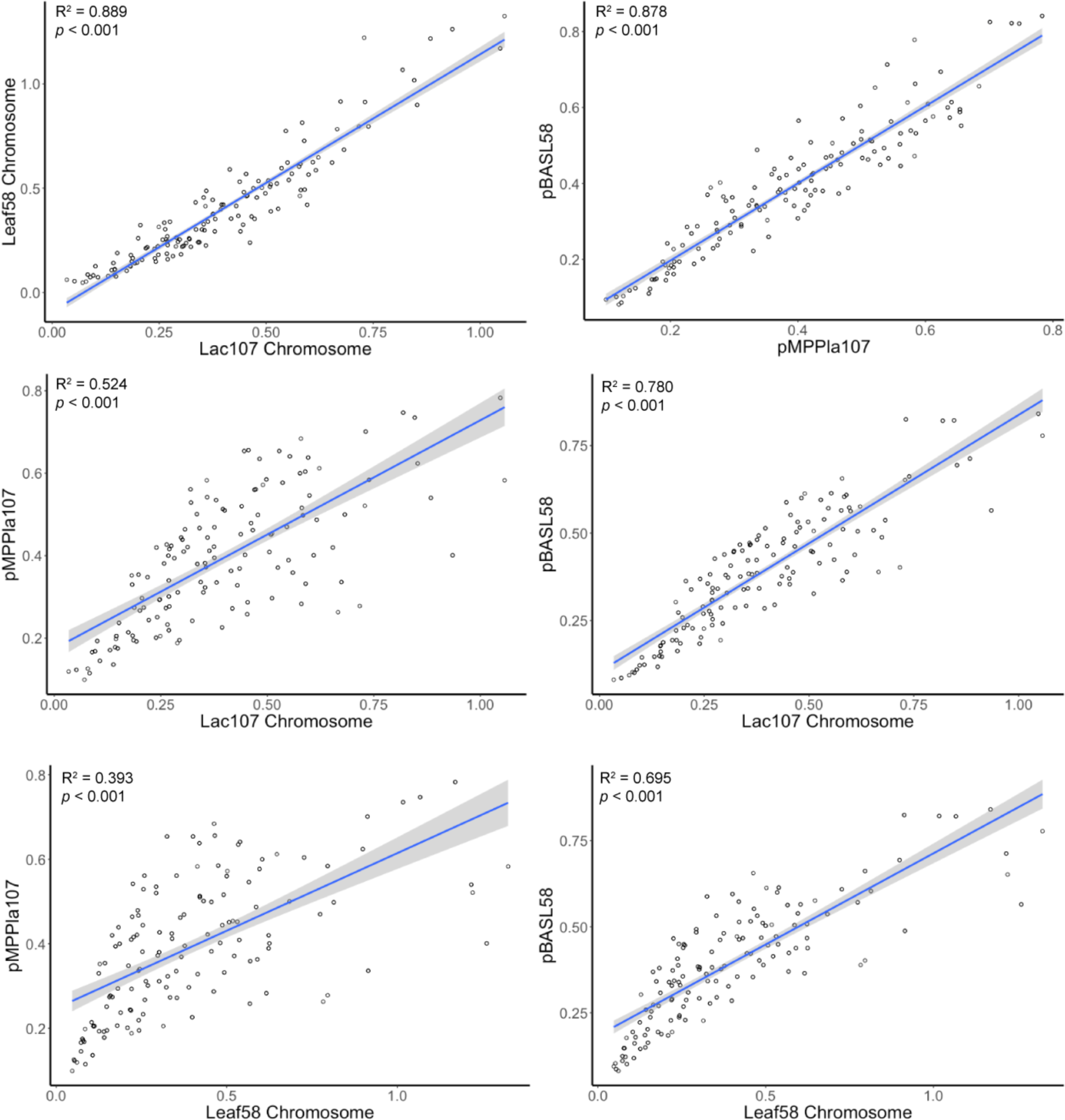
Tetranucleotide frequencies between pMPPla107 and pBASL58 suggest an evolutionary relationship. Nucleotide biases were determined to demonstrate relatedness of megaplasmid and chromosomal sequences in a pairwise fashion. The blue line represents the linear regression model with the surrounding shaded grey area indicating a 95% confidence interval. R^2^ and *p* values are listed for each comparison.

### Housekeeping gene functionality is shared by pBASL58 and pMPPla107

Based on the structural similarities established, we hypothesized that pMPPla107 and pBASL58 would share similar functional pathways. UProC called 9% (85) and 10% (111) of the predicted coding regions for pBASL58 and pMPPla107, respectively, indicating the majority of predicted gene functionality is unknown. Annotation with Prokka returned similar results (13% of genes with annotated functions). The pathways and functions most frequently annotated were replication and repair at 2.3% (22 genes) for pBASL58 and 2.4% (26) for pMPPla107, global and overview maps at 2.1% (20) for pBASL58 and 2.2% (24) times for pMPPla107, and nucleotide metabolism at 1.3% (12) for pBASL58 and 1.8% (19) for pMPPla107 (Figure 5). KEGG KASS also predicated that the two megaplasmids share 57.6% (99/172) of annotated genes. Therefore pBASL58 and pMPPla107 carry 31 and 42 unique genes respectively. Again, the overall distribution of gene products present on both megaplasmids tends towards DNA synthesis, DNA repair, and synthesis of deoxyribonucleotide-triphosphates (Supplemental Table 1 and Supplemental Figure 3). These shared groups include DNA polymerase III subunits, helicases, primase, ligases, recombination proteins, and exonucleases indicating these megaplasmids encode for pathways associated with their maintenance. Other gene products on these megaplasmids are involved in metabolic pathways such as fatty acid biosynthesis, RNA degradation, Aminoacyl tRNA biosynthesis, and NOD-like receptor signaling pathways. Interestingly, both plasmids also encode for several membrane and multidrug efflux pump genes. Both shared efflux genes belong to the Resistance-Nodulation-Division (RND) family of transporters and are known for their multidrug resistance efflux capabilities indicating potential selective factors enabling maintenance in host cells.

**Figure 5:**
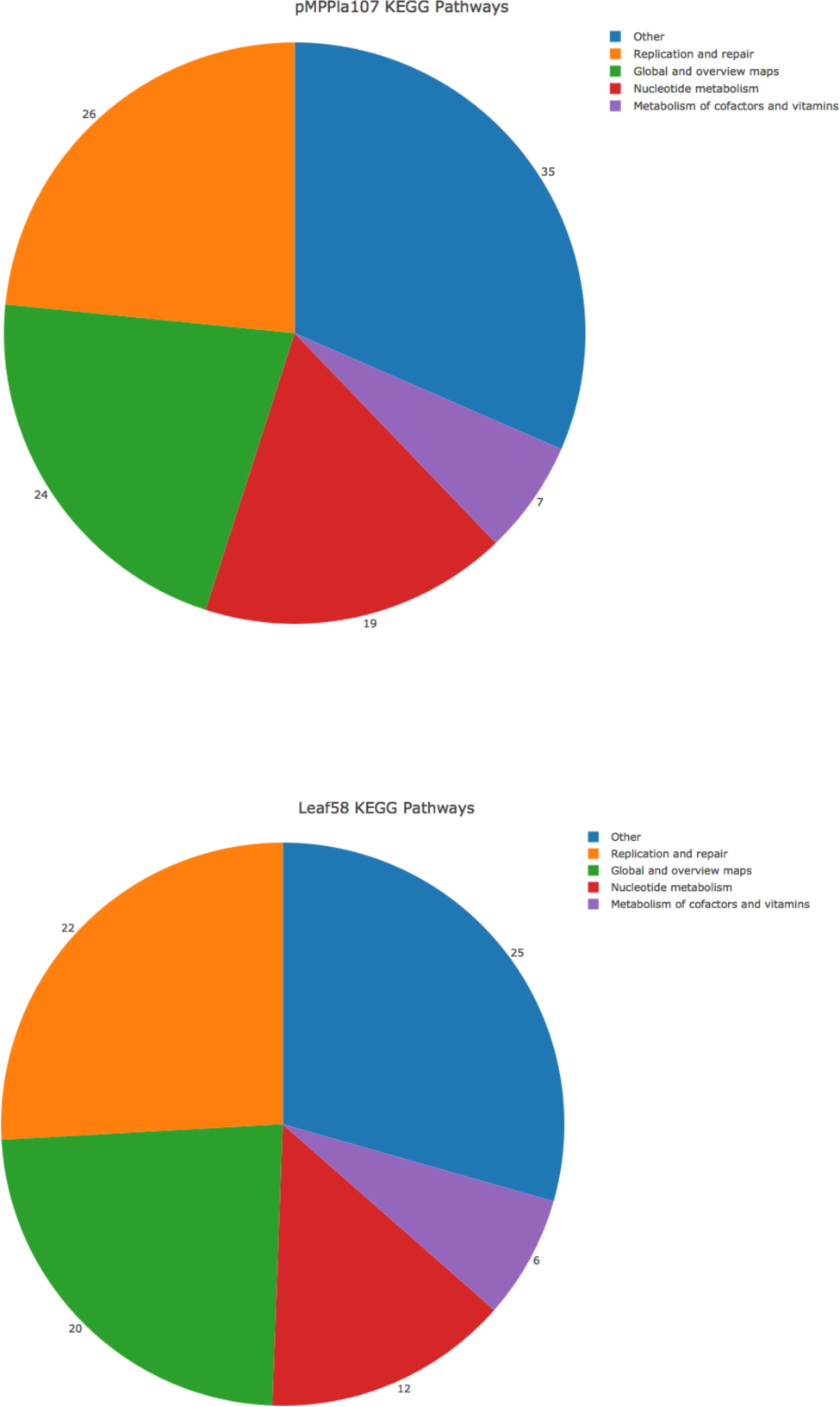
pMPPla107 and pBASL58 share similar functional profiles. pMPPla107 and pBASL58 were input into UProC which counts the number of amino acid sequences that are predicted to belong to a KEGG pathway IDs. KEGG IDs were then matched with the correct pathway. All functional groups with < 5 counts were grouped into an “other” category.

### Differences of pMPPla107 and the Leaf58 plasmid

pBASL58 is predicted to encode a complete CRISPR system from 229-241kb, including two *cas*, three *csy* genes, and a repeat region that includes 36 repeats and spacers(Figure 6). This CRISPR is located in the region of dissimilarity between pMPPla107 and pBASL58 and is not found in pMPPla107. pBASL58 and pMPPla107 do share a (presumably) incomplete CRISPR systems at 436kb and 576kb respectively (Figure 6). These regions include *cas3, csy3*, and *csy4* but lack *csy1, csy2.* pMPPla107 lacks a repeat region altogether associated with this locus while pBASL58 has a repeat region at 720kb encoding 9 repeats and spacers. To our knowledge these are the first complete CRISPR systems located on plasmids found within Proteobacteria.

**Figure 6:**
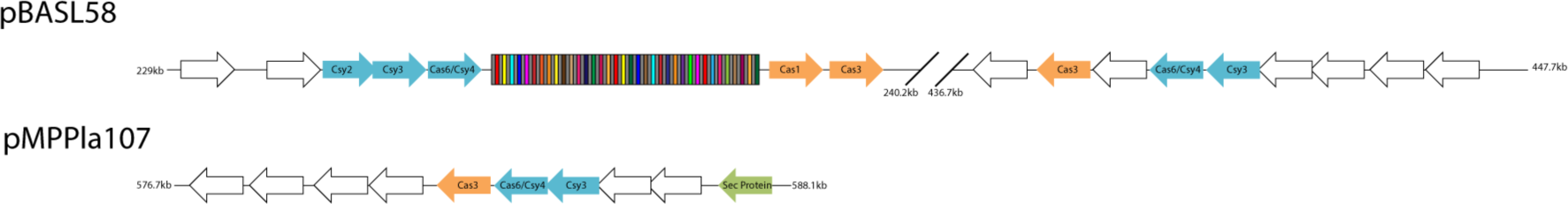
CRISPR systems on pBASL58 and pMPPla107. pBASL58 encodes two CRISPR loci, one of which contains a repeat-spacer regions of 36 repeats. pMPPla107 contains a CRISPR locus without any repeat-spacer regions. Direction of arrows indicates gene orientation. Arrows are colored as: blue) *cys* genes, orange) *cas* genes, green) secretion genes, and white) hypothetical genes. The multicolored boxes indicate the repeat-spacer region, where grey boxes are spacers and colored boxes are repeats.

There exists a region of dissimilarity across both megaplasmids, occurring after approximately 170kb (Figure 3), which could be classified as a cargo region. In pMPPla107 this region consists of 468 predicted genes, of which 27 are annotated. 18 of these 27 annotated genes can be found in pBASL58 and again encode for genes associated with DNA replication, repair, and metabolism. These genes also include membranous proteins like FtsH, which is known to degrade unnecessary or damaged membrane proteins ^45,46^. We have also found that this region can largely be deleted from pMPPla107 during lab adaptation (unpublished) even though the rest of the plasmid is maintained. These data suggest that although this large region may be expendable in some strains, pBASL58 has maintained many of the annotated genes perhaps pointing to their importance in megaplasmid stability or maintenance.

## Discussion

We report a family of divergent, yet syntenic megaplasmids found in single isolates across distinct *Pseudomonas* species. High levels of synteny are matched by shared signals in both tetranucleotide bias and protein pathway functionality. However, these plasmids hosted by strains that are phylogenetically and geographically separated; Pla107 (containing pMPPla107) was found within a *P. syringae* isolate as a causative agent of cucumber disease in Japan, while Leaf58 was found as an epiphyte of *Arabidopsis* in Switzerland in a strain most closely related to *P. putida^41^.* Furthermore, despite high levels of synteny and shared protein functionality, consistently high levels of divergence across shared proteins (≈30%) suggest both plasmids have been independently evolving for a relatively long period of time. From this data we infer that multiple additional members of a family of relatively large (≈1Mb) “cryptic” megaplasmids likely persist within *Pseudomonas* strains.

That there have been no signs of these megaplasmids in the numerous sequences of pseudomonads closely related to each of these isolates is strong indication that these megaplasmids have been relatively recently acquired by their host strains. This pattern, coupled with high levels of divergence between members of this megaplasmid family, suggest that these replicons likely have a high turnover rate within strains over evolutionary time and may persist within communities through frequent horizontal transfer. In other words, presence of this megaplasmid family may be transient in any given genome, but has likely been maintained within Pseudomonads for a long time. Such a lifestyle is consistent with high levels of conjugation as observed in pMPPla107 under laboratory conditions^25^.

Replication, transcription, and translation of horizontally transferred genes are known to incur costs on host cell resources with protein production likely having the greatest effect on fitness^15,47-49^. Previous work on pMPPla107 suggests that acquisition of the megaplasmid results in lowered fitness and other phenotypic changes which could be costly in some environments, yet it still transfers readily and is maintained within host cells^25,26^. Such costs could likely be the reason pMPPla107 and pBASL58 encode a large number of genes involved in critical functions regarding plasmid maintenance and transmission as well as potential addiction systems and could enable long-term survival despite a transient lifestyle. In particular, there are various proteins found in pMPPla107 and pBASL58 involved in synthesizing precursors for nucleotides such as: thymidylate synthase, guanylate kinase, ribonucleoside diphosphate reductase, deoxycytidine triphostphate deaminase, and glutamate synthase (Supplemental Table 1 and Supplemental Figure 3). The megaplasmids may carry these proteins in order to increase flux to nucleotide synthesis and drive replication and transcription processes to alleviate any physiological costs an additional ≈1Mb of newly acquired DNA may bring. Many of these genes do not encode for complete pathways, indicating possible parasitic behavior of host resources while ensuring the necessary building blocks for plasmid maintenance are available.

Plasmid usage of host tRNA pools has been shown to deplete tRNAs resulting in reduced growth and fitness^20,50-52^. The large number of tRNAs and presence of a handful of annotated tRNA ligases encoded on the megaplasmids may serve the purpose of avoiding translational costs due to tRNA depletion or may accommodate codon usage bias between chromosome and megaplasmid. Both megaplasmids are also predicted to encode Mfd, Rep, DnaB, and RecA all known to resolve replication and transcription complex conflicts ensuring successful replicon duplication and transcription^53,54^. We hypothesize the megaplasmids maximize their ability to persist by eliminating or compensating for these potential costs by encoding a variety of housekeeping genes coupled with high levels of horizontal transfer through conjugation.

Evolutionary relationships between pBASL58 and pMPPla107, their relatively large size and contribution to gene content of single strains, coupled with maintenance of “housekeeping” genes, and high levels of transfer across pseudomonads suggest that this megaplasmid will provide unique insights into an evolutionary argument concerning horizontal transfer referred to as the complexity hypothesis^17^. The complexity hypothesis has been through multiple revisions, but is currently interpreted as a trend where horizontally transferred genes are less likely to be involved with complex processes (like translation) and maintain a lower number of protein-protein interactions than vertically inherited loci^4^. One current limitation of the complexity hypothesis, as highlighted by these megaplasmid families, is that is fails to reconcile gene conservation in the context of highly mobile selfish DNA like plasmids. Both pBASL58 and pMPPa107 contain numerous “complex” genes, including those involved in nucleotide synthesis, DNA replication, and translation and yet these genes are clearly horizontally transferred across strains. Therefore, the presented family of megaplasmids potentially necessitates a caveat to the complexity hypothesis in which “complex” genes can be horizontally transferred frequently but aren’t maintained over time, because they are linked together on megaplasmids that require these pathways to ameliorate physiological costs.

Likewise, there have been numerous recent discussions about whether bacterial pangenomes are adaptive or neutral. Similar to the complexity hypothesis, these discussions tend to focus on the presence/absence of single genes across a variety of closely related genomes rather than the linked gain/loss of genes that compose a pangenome^1-3^. To put this in perspective, recent findings suggest that the *P. syringae* pangenome is composed of 77,728 genes, meaning that 1.5% of these are solely present on pMPPl107^55^. Since megaplasmids have the potential to add thousands of genes to a pangenome linked together in a single transfer event^56^, one has to consider that evolutionary pressures may act differentially on subsets of the pangenome. Our data suggest that a majority of genes on these megaplasmids may be either neutral or costly to the host when selection is considered in the context of the host genome. However, a majority of genes linked on the megaplasmid may be selectively beneficial for megaplasmid maintenance and/or transfer regardless of fitness of the host cell. Thus, presence of a majority of genes on the megaplasmid (and which are part of the pangenome) are under selection at some level, but only a minority of these may be beneficial at the level of bacterial strains or populations.

CRISPR-Cas systems have become popularized recently because of their utility in genome editing, however, these systems likely originated in bacteria as defense mechanisms against invasion of foreign genetic material^57-61^. CRISPR arrays are often carried and transferred by larger plasmids in bacteria and archaea, yet *cas* genes are rarely found on plasmids^62,63^. Here we characterize a potentially shared CRISPR-Cas system bound to the bacterial megaplasmids pMPPla107 and pBASL58. Although pBASL58 encodes a fully intact CRISPR-Cas3 system with a region containing 36 spacers and repeats, this repeat and spacer region are not present within pMPPla107 leading us to believe pMPPla107’s system is nonfunctional. Regardless of functionality, it is quite interesting that at least one of these megaplasmids contains an intact CRISPR locus given the widespread idea that these systems are used by bacteria to defend against parasites and mobile elements. Perhaps the presence of a CRISPR system is a beneficial and selective trait for retention of pBASL58 in host cells in that it provides a transferable immune pathway. However, the recent description of CRISPR spacers that target sites on bacterial chromosomes also suggest that these loci may also function in gene regulation ^64-67^.

Using comparative computational and molecular approaches we have characterized pBASL58, the second member of a family of large megaplasmids found in Pseudomonads. Conservation of pathway presence and megaplasmid structure strongly suggests that a majority of the sequences on pBASL58 and pMPPla107 have diverged from a common ancestral plasmid. However, the consistent levels of divergence between proteins shared by both plasmids suggest that this common ancestral plasmid did not recently exist. Finding two related plasmids with such high level of divergence also highlights the likelihood that other members of this megaplasmid family exist in nature and are waiting to be found. Our work serves as a guide to discover megaplasmid families as well as a foundation of understanding the forces that structure megaplasmid evolution, maintenance, and transfer.

## Supplemental Figures

**Supplemental Figure 1.**
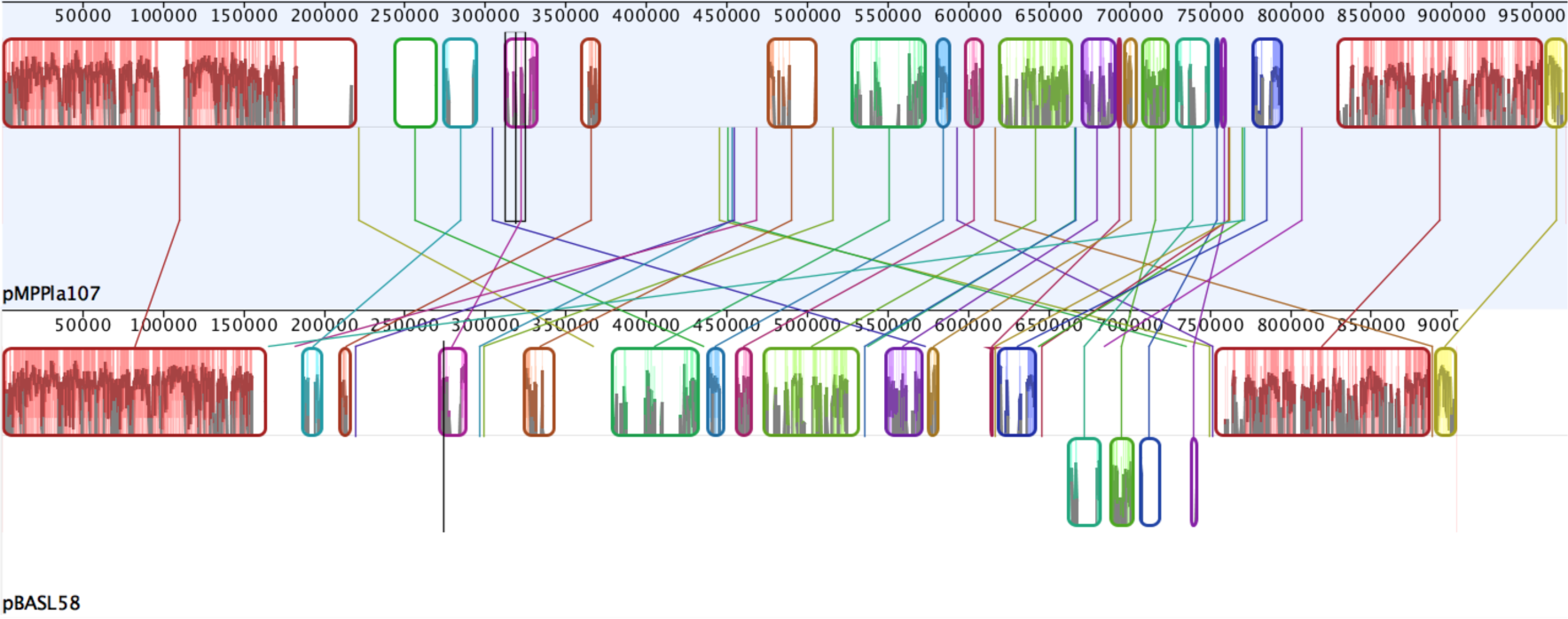
MAUVE analysis of pMPPla107 and pBASL58 demonstrate local collinear blocks (LCBs) and areas of synteny. Lines connect LCBs with each other between megaplasmids. Blocks below the midline for each sequence indicate inverted regions. Colored areas within LCBs indicate higher levels of homology between sequences.

**Supplemental Figure 2.**
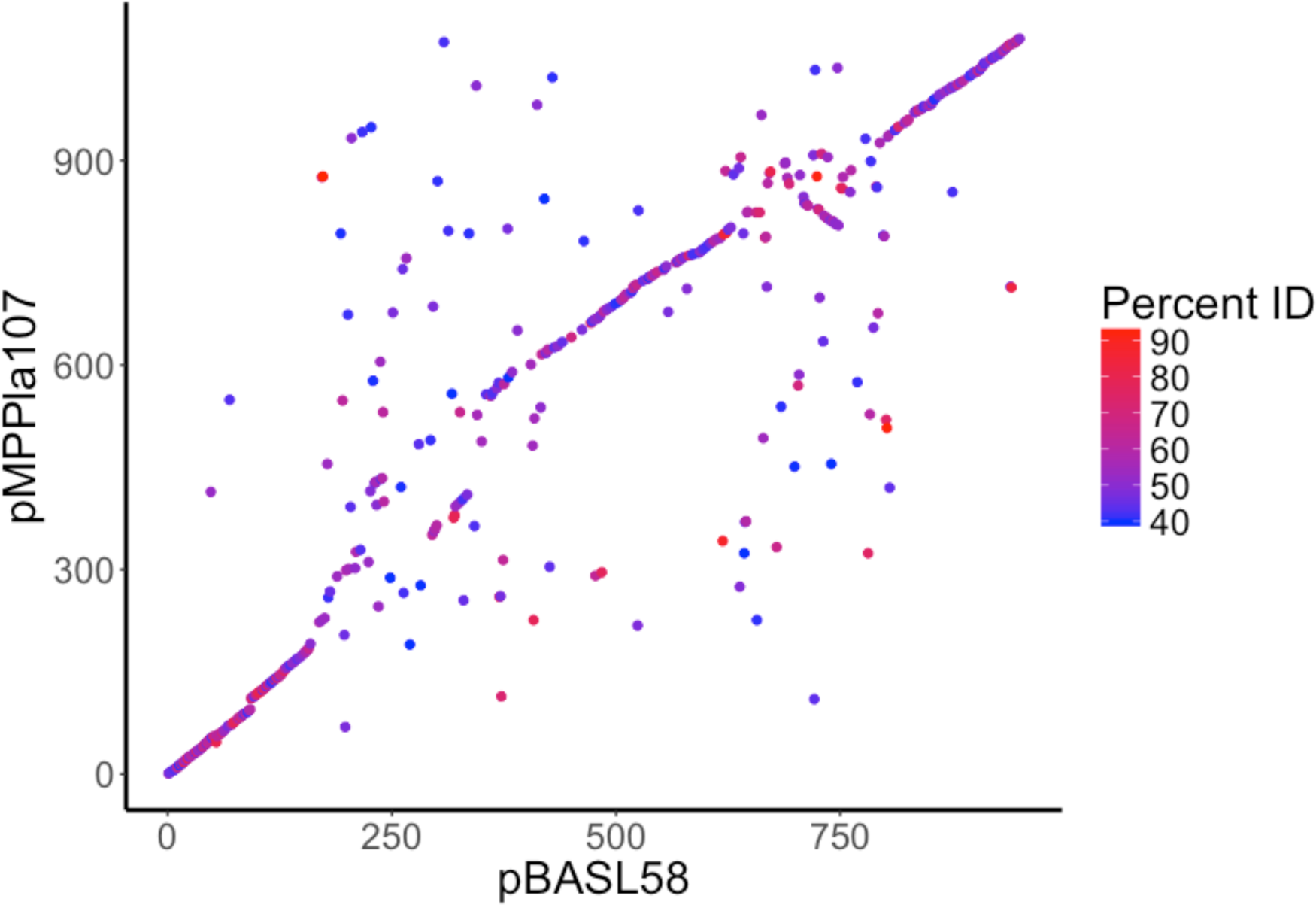
BLASTp results of pMPPla107 and pBASL58 indicate syntenic and divergent sequences. Axes indicate gene position order. The color gradient is set to BLASTp percent identity results for each comparison. A percent identity cutoff of ≥40% was used.

**Supplemental Figure 3.**
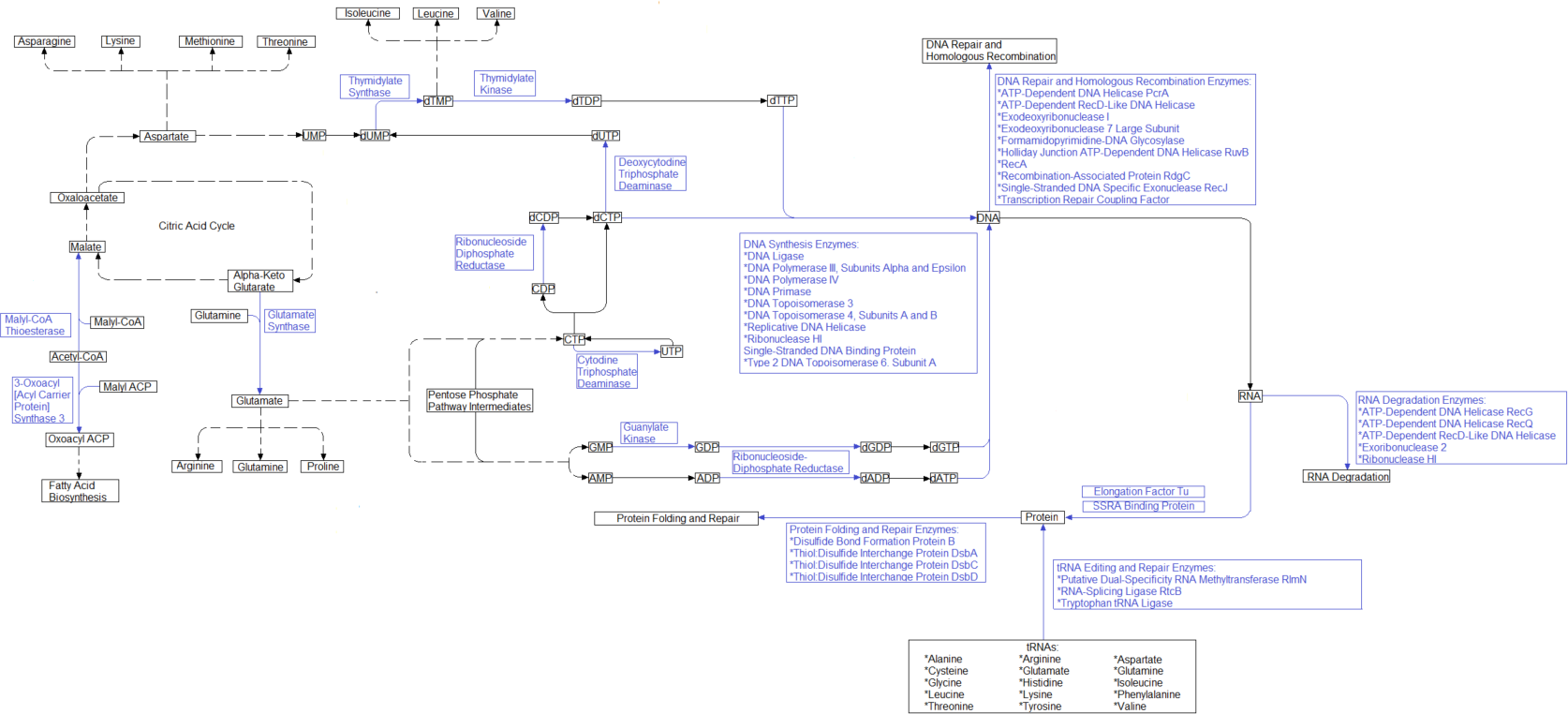
Metabolic pathways shared by pBASL58 and pMPPla107. Output of KEGG/KASS pathways between pBASL58 and pMPPla107 were matched and are reported by blue pathways with their associated genes. Hashed lines indicate that no genes were found in pBASL58 or pMPPla107 to be involved with that metabolic process.

